# Positive interactions within and between populations decrease the likelihood of evolutionary rescue

**DOI:** 10.1101/2020.08.06.239608

**Authors:** Yaron Goldberg, Jonathan Friedman

## Abstract

Positive interactions, including intraspecies cooperation and interspecies mutualisms, play crucial roles in shaping the structure and function of many ecosystems, ranging from plant communities to the human microbiome. While the evolutionary forces that form and maintain positive interactions have been investigated extensively, the influence of positive interactions on the ability of species to adapt to new environments is still poorly understood. Here, we use numerical simulations and theoretical analyses to study how positive interactions impact the likelihood that populations survive after an environment deteriorates, such that survival in the new environment requires quick adaptation via the rise of new mutants - a scenario known as evolutionary rescue. We find that the probability of evolutionary rescue in populations engaged in positive interactions is reduced significantly. In cooperating populations, this reduction is largely due to the fact that survival may require at least a minimal number of individuals, meaning that adapted mutants must arise and spread before the population declines below this threshold. In mutualistic populations, the rescue probability is decreased further due to two additional effects - the need for both mutualistic partners to adapt to the new environment, and competition between the two species. Finally, we show that the presence of cheaters reduces the likelihood of evolutionary rescue even further, making it extremely unlikely. These results indicate that while positive interactions may be beneficial in stable environments, they can hinder adaptation to changing environments and thereby elevate the risk of population collapse. Furthermore, these results may hint at the selective pressures that drove co-dependent unicellular species to form more adaptable organisms able to differentiate into multiple phenotypes, including multicellular life.

## Introduction

Positive interactions play key roles in shaping the assembly, function and evolution of many ecological communities^1–3^. Extensive research has demonstrated the prevalence of positive interactions in numerous ecosystems, ranging from plant communities to the human microbiome^4–9^. Positive interactions occur both as intraspecies cooperation, such as bacterial populations that are able to resist antibiotics by collectively secreting antibiotic-degrading enzymes^10,11^, and as interspecies mutualism, such as the cross-protection relationship between sea anemones and clownfish^12^.

Evolutionary theory demonstrates that positive interactions can be selected for. For instance, positive interactions such as nutrient exchanges between individuals, can arise due to the benefit of removing costly genes required for the generation or acquisition of the exchanged nutrients. This idea, termed ‘The black queen hypothesis’, is proposed as a dominant force that promotes an increase in the abundance of positive interactions between species^13,14^. Positive interactions may also be beneficial for the entire population through ‘division of labour’ - a situation where individuals exchange the products of different tasks in which they specialized and can perform efficiently^15^. Such benefits of positive interactions have been implicated in the evolution of multicellularity, owing to the resemblance of multicellular organisms to unicellular species that form genetically identical subpopulations of cells with different phenotypes that attain division of labour^16–18^. A well-known example of division of labor is the filamentous photosynthetic cyanobacteria that form subpopulations of nitrogen-fixing heterocysts that enable different cells to exchange the benefits of nitrogen-fixation and photosynthesis^19^. Thus, insights regarding the evolutionary dynamics of positive interaction might shed light on the transition from unicellular to multicellular life.

While the formation of positive interactions may be selected for, populations engaged in positive interactions can have heightened sensitivity to biotic and abiotic stresses. When species are interdependent, a stress that affects one species may have cascading effects that lead to the extinction of multiple additional species. This phenomenon, termed co-extinction, is found in many conservation studies examining the effects of anthropogenic environmental changes^20–22^. In addition, cooperating populations are prone to invasion by non-cooperating ‘cheaters’ that spread at the expense of the cooperators and can even lead to their collapse^23,24^. Such collapses occur since interactions are typically multifaceted: individuals may cooperate in one task, and simultaneously compete in another. For example, populations containing antibiotics degrading bacteria may collapse due to the rise of non-degrading cheaters that are able to outcompete the degraders for nutrients^25^ More broadly, positive interactions and competition often occur concomitantly and form a cooperation - competition continuum^26–28^. The unstable nature of positive interaction likely has significant effects on the evolution of populations and communities, but thus far research in this area has focused primarily on the sensitivity of cooperating populations to cheaters^29–31^.

In particular, the effect of positive interactions on adaptation to changing abiotic environments is still poorly understood. Several recent studies suggest that adaptation may be hindered by positive interactions. First, the response of ecosystems to climate change suggests that mutualisms are unstable when adapting to novel environments^32,33^. A notable example is the cross protection relationship between sea anemones and clownfish, that was shown to be perturbed by the imbalance of their adaptation rate^32^. Next, several experimental evolution studies involving bacterial mutualisms have found that while some populations evolve a more efficient division of labour and elevated growth rates, other replicate populations experience unexplained collapse, leading to extinction of both interacting species^34,35^. Finally, a recent study exploring the adaptability of metabolically co-dependent bacterial populations in the presence of antibiotics showed that codependency between two species results in a limitation of their adaptation ability by the least adaptable, “weakest link” strain^36^. Thus, once a species is more adapted to the environment than its partner, it can not further increase in fitness until it’s cooperator evolves to a similar fitness. Taken together, these findings suggest that limited capacity for adaptation may be a general implication of positive interactions, but we still do not have a clear understanding of the severity of the limitations and the mechanisms causing it.

To address these questions, we focus on the extreme case of adaptation following a deterioration of a formerly hospitable environment into an inhospitable one, in which a population or community is heading towards extinction (**Fig. 1A**). In this scenario, survival requires rapid adaptation to the new environment, which is possible only through the rise of new mutants - a phenomenon termed “Evolutionary rescue”^37,38^. When populations do not engage in positive interactions, it is sufficient for adapted mutants to arise prior to the ancestor’s extinction in order to rescue the population (**Fig. 1B**). We elucidate how different types of positive interactions influence the likelihood of evolutionary rescue by conducting numerical simulations and theoretical analyses. We conclude that while positive interactions may be beneficial in steady environments, they can hinder adaptation to changing environments. These results may offer new insights into the evolutionary dynamics of ecosystems facing sudden environmental stress, such as climate change, and the selective forces that affect cooperative and mutualistic populations over evolutionary time scales.

**Figure 1:**
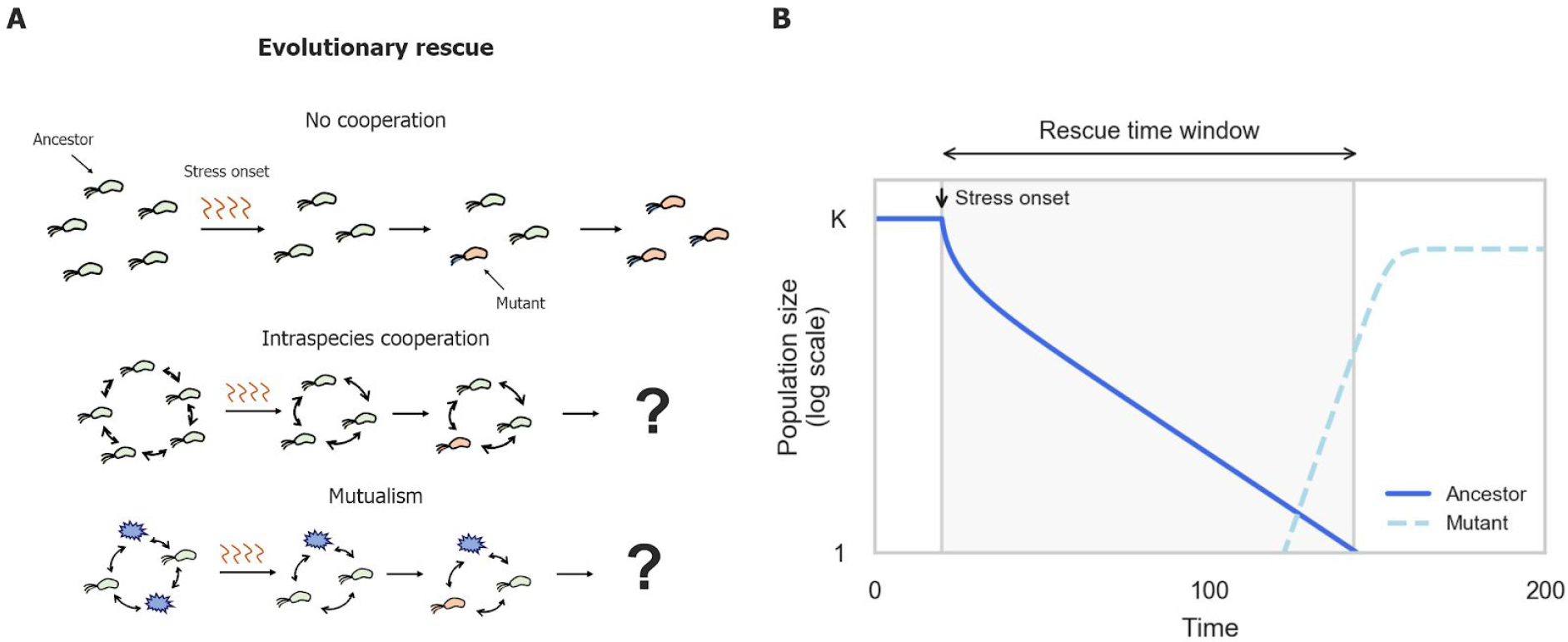
Evolutionary rescue in populations engaged in positive interactions is poorly understood. (A) An abrupt stress in the environment causes population density to decline toward extinction (green and purple cells), followed by adaptation via the rise of new mutants (orange cells). (B) Example simulation of evolutionary rescue in a non-cooperating population. Rescue time window, the time window during which adapted mutants can rise and prevent the population’s extinction, is calculated as the time it takes the ancestors to become extinct. In our simulations, a population is considered extinct when it reaches below 1 individual. Parameter values used in simulations are provided in Table S1.

## Results

### Cooperative populations have a limited time window for evolutionary rescue

In order to analyze the dynamics of intraspecies cooperation when evolutionary rescue is required, we have added an evolutionary component to a previously established ecological model of cooperative populations^40^ (**Fig. 1**, **Eqs. 1–3**, Methods, and Supplementary Information). Briefly, the model is based on the classical logistic growth model, in which populations initially grow at rate (*r*) and saturate at carrying capacity (*K*). It extends the logistic model by reducing individuals’ growth rate when the population is below a critical size (*N_c_*), a phenomenon known as an Allee effect’^39^ (**Fig. S1**). For example, bacterial populations experience an Allee effect when collectively degrading antibiotics since their growth rate is increased only if there are enough degrading cells to sufficiently reduce the concentration of antibiotics in the environment. In addition, we included a death rate (*δ*) reflecting an external environmental stress that is independent of the interactions within the populations. In the example of bacteria collectively degrading antibiotics, such a stress may be a rise in temperature that impairs the bacteria’s growth without affecting the antibiotic. Our model includes an ancestral population (*A*) and a mutant population (*M*) with an increased growth rate (*r_A_* < *r_M_*). The dynamics of the ancestor and mutant populations are given by:

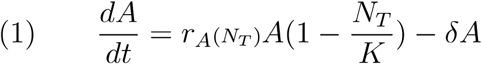

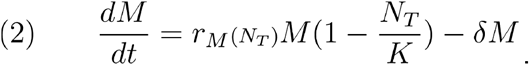

The growth rate *r* depends on the total population size (*N_T_* = *A* + *M*):

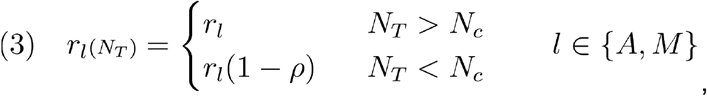

where *ρ* ∈ [0, 1] is the fraction by which growth rates decrease below the critical population size (*N_c_*). The strong Allee effect was implemented as a step function in order to enable analytical calculations and maintain simplicity. Qualitatively similar results also occur when the Allee effect is modeled using a more complex, smooth function (**Fig. S2,3** and Section 4 in the supplementary Information).

The evolutionary rescue scenario was explored by running simulations in which an ancestor population experiences an abrupt increase in environmental death rate which causes the population to decline toward extinction, and mutants may arise stochastically during this decline, potentially spreading and rescuing the population from extinction (Fig. 1B). Each simulation begins with the growth of an ancestor population in an unstressed environment (*δ* = 0), followed by an onset of stress that increases the death rate such that it exceeds the ancestral exponential growth rate (*r_A_* < *δ*), leading the population to decline toward extinction (Fig. S1A). Mutation events are modeled as a poisson process, with the expected number of mutants at each time interval given by the ancestral population size and mutation rate (μ) (**Eq. S4**). Mutants differ from ancestors only in their elevated exponential growth rate (*r_A_ < r_M_*), which allows them to survive the stress, but only when the total population size exceeds the critical threshold (*r_M_* > *δ* > *r_M_* – *ρ*) (**Fig. S1B**). For simplicity, no further stochastic effects were considered in this model. We assessed the evolutionary rescue probability by calculating the fraction of simulations in which the mutants were able to spread and exceed the critical population size.

Consistent with previous works^40^, we found that populations engaged in intraspecies cooperation have lower probability of evolutionary rescue in comparison to non-cooperative populations (**Fig. 2A, S8**). A main cause of this reduced rescue probability is that in cooperative populations adapted mutants can spread only if they appear while the total population size exceeds the critical size (**Fig. 1B**), while in non-cooperative populations adapted mutants can spread regardless of the total population size. Thus, the critical population size limits the rescue time window - the time window during which adapted mutants can rise and prevent the population’s extinction (**Fig. 2B-C**). The reduction in rescue probability occurs even when cooperation provides a fitness advantage. Cooperative populations only have a rescue probability comparable to that of non-cooperative ones when their growth rate is significantly higher - up to twice that of non-cooperating populations for large critical population sizes (**Fig. S6**).

**Figure 2:**
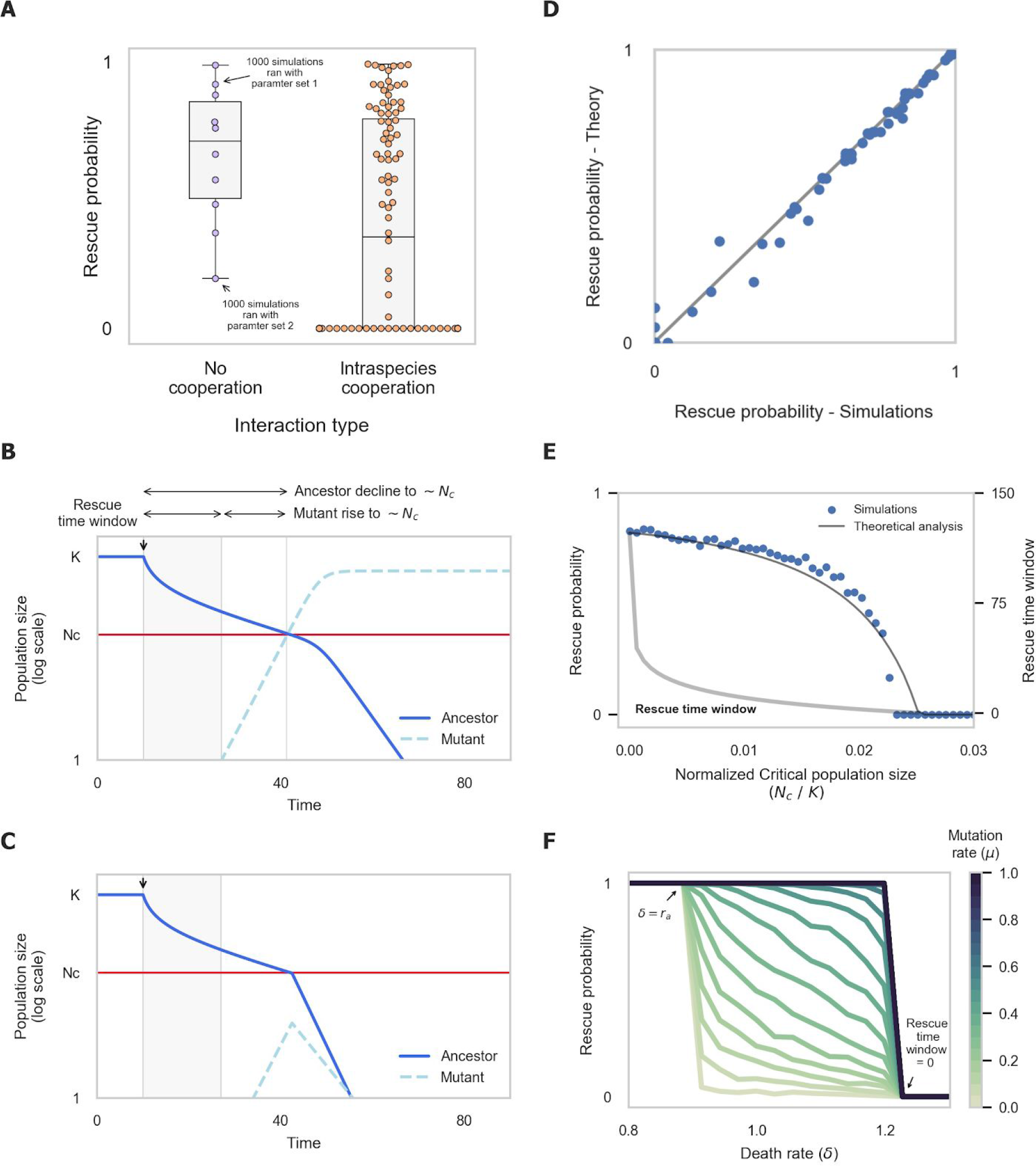
Cooperative populations have a limited time window for evolutionary rescue. **(A)** Intraspecies cooperation has lower rescue probability in comparison to populations with no positive interactions. Each dot represents the rescue probability resulted from 1000 simulations run with different set of parameters (Critical populations size (*N_c_*), ancestor and mutant’s growth rate (*r_a_, r_m_*)). (B + C) Evolutionary rescue of populations engaged in intraspecies cooperation requires the mutant’s population to reach critical population size before ancestor. This results in a shorter rescue time window, outside of which mutants cannot reach critical population size and rescue the population. (D) Theoretical analysis matches well the rescue probability observed in simulations. (E) The rescue probability and rescue time window decrease as the critical population size increases. (F) Death rate (*δ*) and mutation rate (*μ*) effect on evolutionary rescue. Mutation rate sets the sharpness of the transition between certain rescue (*P* = 1) and certain extinction (*P* = 0).

The rescue time window and rescue probability decrease as the critical population size increases. An analytical approximation of the rescue time window is given by the difference between the time it takes the ancestral population to decline to the critical population size, and the time it takes adapted mutants to grow sufficiently (Section 3 in the supplementary Information). The probability that an adapted mutant arises during this time interval provides an excellent approximation of the rescue probability observed in simulations (**Fig. 2D**). Notably, when the critical population size is too large, mutants are unable to reach it even if they appear immediately following the onset of the stress (**Fig. 2E**). Thus, in such populations the stress inevitably leads to extinction, unless adapted mutants are already prevalent enough in the population prior to the stress’ onset. The mutation rate affects the likelihood of rescue when the rescue time window exists, but rescue is not possible for any mutation rate when the rescue time window is zero (**Fig. 2F**). These results suggest that the higher the number of individuals required for successful cooperation, the lower the probability to adapt.

### Mutualisms have a greatly reduced probability of evolutionary rescue

Next, we test how mutualistic interactions affect the probability of evolutionary rescue. To do so, we have used an extended model in which two species are dependent on each other in an obligatory manner (**Eqs. 4–6**, Methods and Supplementary Information). Analogously to the case of intraspecies cooperation, the growth rate of each species is reduced when the population size of its partner is below a critical population size (*N_c_*):

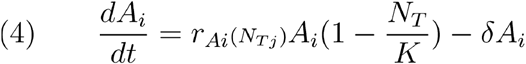

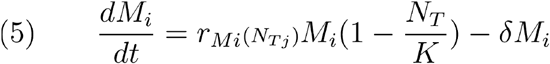

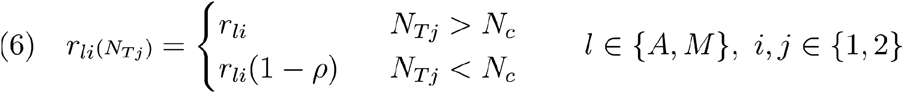

We have again assessed the rescue probability by running simulations over a range of parameters and measuring the fraction that resulted in survival of the two species. Qualitatively similar results also occur when the mutualistic interactions are modeled using a more complex, smooth function (**Fig. S4**) and when the initial densities of the species are unequal (**Fig. S9**).

We observe that the probability of evolutionary rescue of populations engaged in mutualistic interactions is significantly lower than that of cooperative populations (**Fig. 3A**). Since mutualistic interaction can provide fitness advantage through division of labor, we have also compared the rescue probability of populations engaged in intraspecies cooperation with that of mutualistic populations that have a higher growth rate. We found that the growth rates of mutualistic populations must be greater by up to 30 percent in order for their rescue probability to be equal to that of cooperating ones (**Fig. S7**).

**Figure 3:**
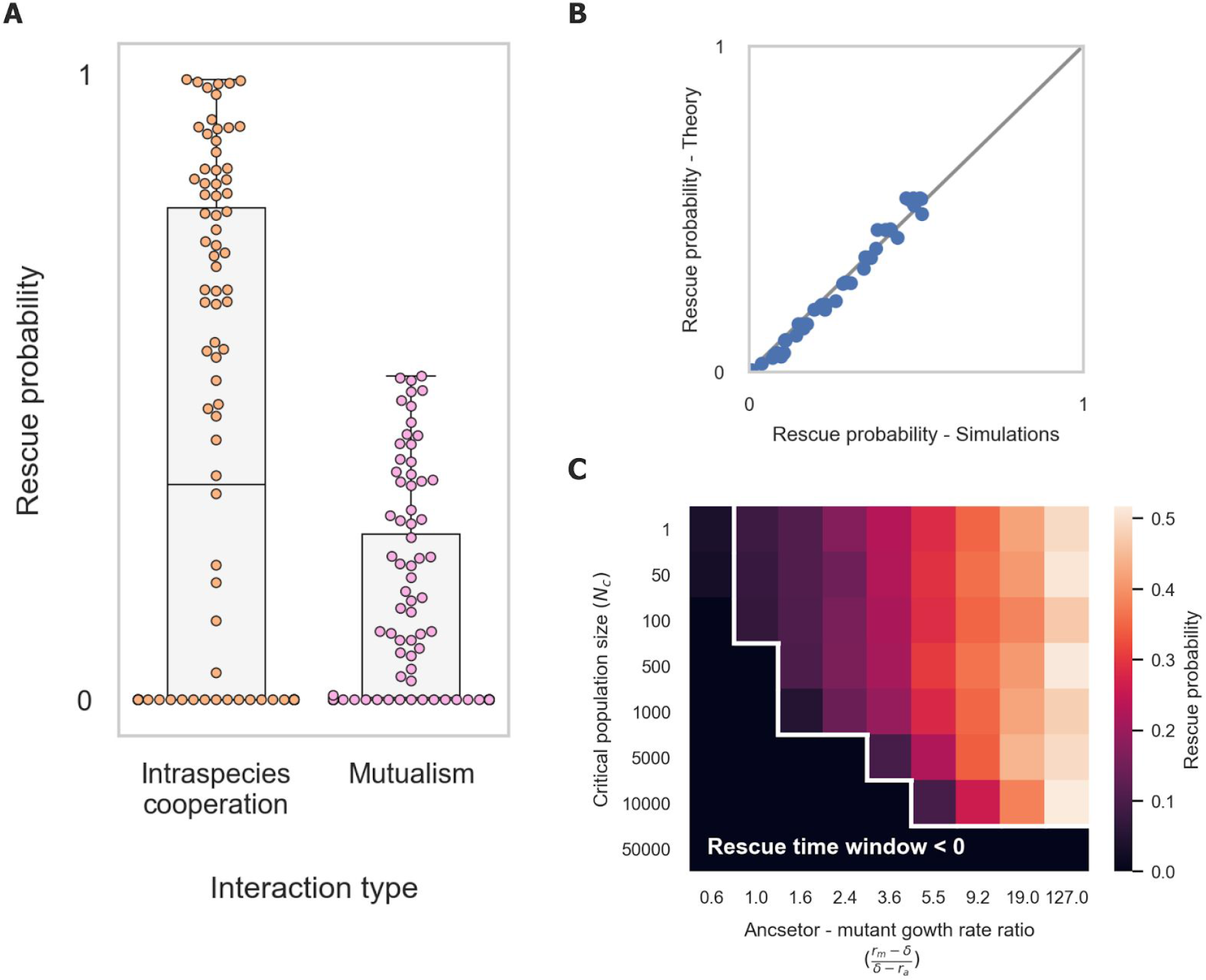
Mutualisms have a greatly reduced probability of evolutionary rescue. (A) Rescue probability is greatly reduced in mutualisms compared to intraspecies cooperation. Dots represent the rescue probability calculated from simulations run with different sets of parameters as in figure 2. (B) Theoretical analysis matches well the rescue probability observed in simulations. (C) Rescue probability decreases with critical population size (*N_c_*) and the ratio between mutant and ancestor growth rates. As in intraspecies cooperation, the rescue time window reveals a transition curve under which rescue probability is zero.

As in the case of cooperative populations, a theoretical analysis based on the length of the rescue time window approximates the probability of evolutionary rescue well (**Fig. 3B** and Section 3 in the Supplementary Information). We observed again that evolutionary rescue in mutualisms is possible only if adapted mutants arise early enough such that they are able to grow sufficiently before their partners’ population declines below the critical population size (**Fig. 3C**). However, this is not sufficient to explain the lower likelihood of rescue found in mutualisms compared to cooperative populations, as the duration of the rescue time window is identical in both systems.

We found two additional major effects that reduce the rescue probability in populations engaged in under mutualistic interactions (**Fig. 4**). First, since the two species are dependent on each other, adaptation of a single species is not sufficient to rescue it from collapse, even if the adapted mutant arises within the rescue time window (**Fig. 4A-B**). Due to its dependence on the other species’ ability to cooperate, it will collapse as soon as the second species falls below the critical population size. Thus, evolutionary rescue requires adapted mutants to arise and spread in both species in order to rescue either of them, which significantly reduces the evolutionary rescue probability since it requires the occurrence of two independent rare mutation events.

**Figure 4:**
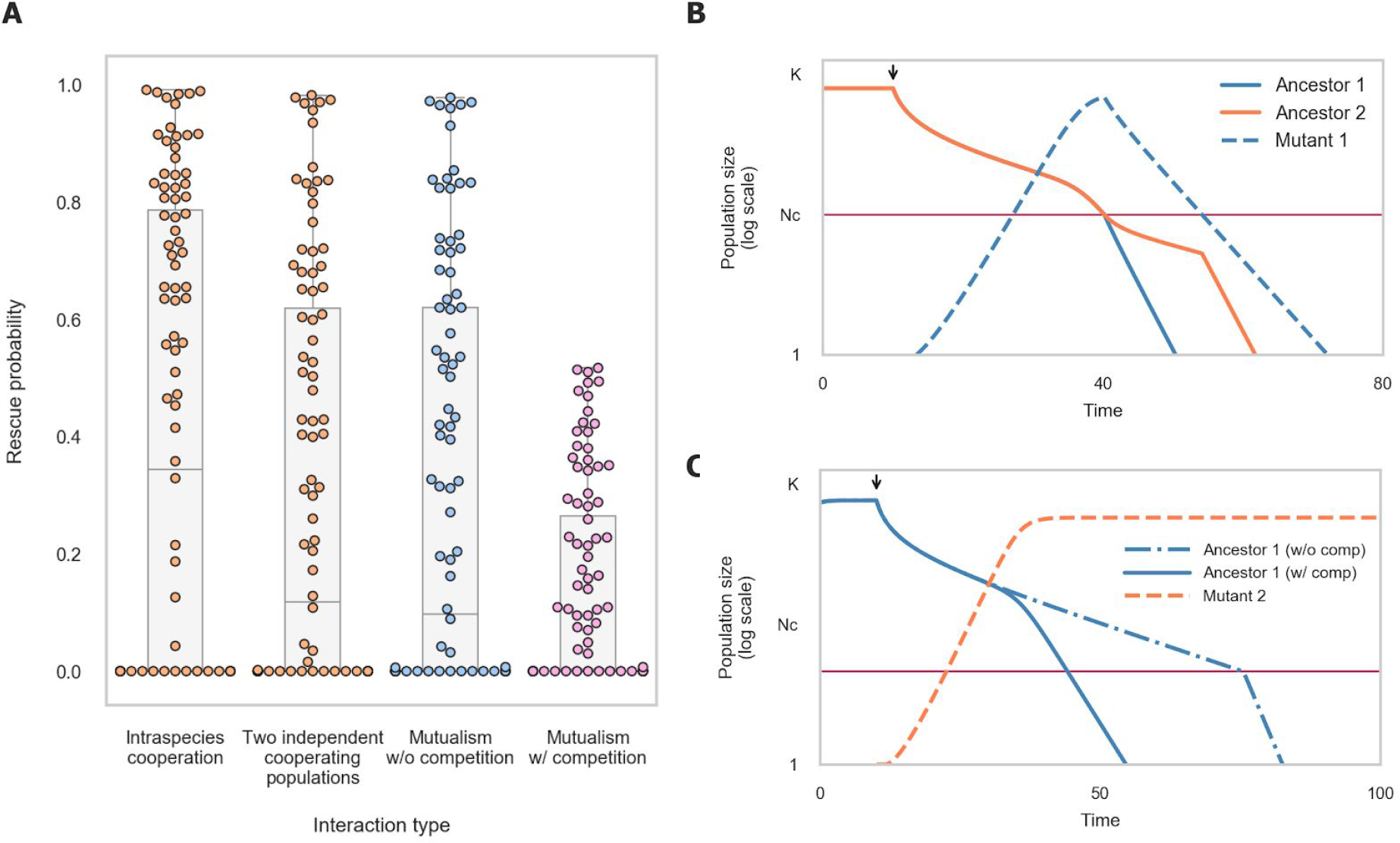
Rescue probability of mutualisms is reduced since it requires adaptation of both mutualistic partners, which also compete for resources. (A) In the absence of interspecies competition, mutualisms have a decreased rescue probability that is similar to that of two independent cooperating populations. Interspecies competition further decreases the probability of rescue of two mutualistic species. Dots represent the rescue probability calculated from simulations run with different sets of parameters as in figure 2. (B) Example of a simulation in which adaptation of a single species within the rescue time window does not suffice to prevent extinction due to its dependence on the other species’ ability to cooperate. (C) Comparison of the rate of decline of a single species (blue) with and without competition. Adaptation of the mutualistic partner (orange) accelerates the species’ decline due to competition for resources. The dynamics of the unadapted partner species are similar with and without competition and are not shown for simplicity.

The evolutionary rescue probability is further reduced when the two species also compete (**Fig. 4A,C**). While the two species facilitate each other’s growth, they may also compete for resources, which becomes the dominant interaction at high population densities. When adapted mutants of one of the species spread and approach the carrying capacity they outcompete their partner species for resources. This results in a faster decline of the partner species toward its critical population size and in a shortened time window for adapted mutants in this species to arise (**Fig. 4C**).

Both the interdependency between species and the competition within mutualism contribute to the decline in their rescue probability. In the absence of interspecies competition mutualisms have a decreased rescue probability that is similar to that of two independent cooperating populations (**Fig. 4A**). Interspecies competition alone decreases the probability of rescue of two non-mutualistic species, but it is the combination of competition and mutualistic interactions that jointly result in the low rescue probability found in mutualisms (**Fig. 4A**). When the two mutualistic species do not compete, their rescue probability can even exceed that of two independent populations engaged in intraspecies cooperation. This occurs since adaptation of one of the species can increase the rescue time window of its partner (**Fig. S10**). However, this phenomena only occurs for a limited set of parameters, and its influence on the rescue probability is relatively small.

These results suggest that mutualism may have a greatly reduced capacity for adaptation. Since different species rely on each other, adaptation of the community requires all partners to adapt. This slows down adaptation and makes it considerably less likely when limited by the supply of adapted mutations. In addition, when mutualistic partners also compete for additional resources, adaptation of one species can hinder other species’ ability to adapt, potentially leading to collapse of the whole community. We conclude that when the environment is unstable, mutualisms may be a fragile and undesirable strategy even when they offer the benefits of division of labour, as the selective pressure on species to quickly adapt to changing conditions may dominate the advantages conferred by gene loss and division of labour.

### In the presence of cheaters, evolutionary rescue of cooperative populations is extremely unlikely

Since cooperating populations are commonly invaded by cheaters, we next explored how the presence of cheaters affects the evolutionary rescue probability of cooperating populations. To do so, we adapted a previously established model that describes cooperators and cheaters dynamics and was shown to successfully capture the dynamics of yeast populations cooperating in extracellular sucrose degradation in the presence of non-degrading cheaters (**Eqs. 7–10**)^41^. In this model, both cooperators and cheaters are affected by the cooperator’s population density, and have a reduced growth rate when the cooperator population is below a critical size. However, cooperators and cheaters coexist since cheaters have a growth advantage (*b*) at high cooperator density since they do not pay the cost of cooperation, whereas at low populations densities cooperators have a growth advantage (*a*) at low populations densities, reflecting their preferential access to the public goods they produce:

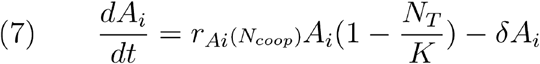

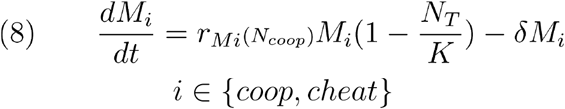

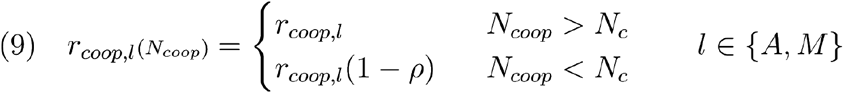

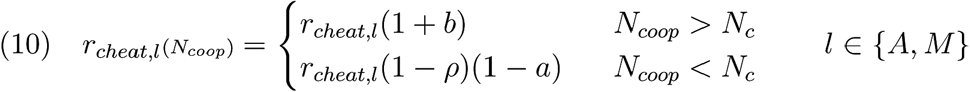

While this model results in oscillatory dynamics, qualitatively similar results are also found in a more complex model in which oscillations do not occur (Fig. S5).

Invasion by cheaters reduces the cooperators’ density to close to the critical population size, which dramatically reduces the rescue time window and the rescue probability (**Fig. 5A**). For the population to survive an adapted cooperator mutant must arise and reach sufficient population size. In contrast, adapted cheaters do not contribute to the population’s growth rate and cannot prevent its extinction. Prior to the stress’s introduction, the presence of cheaters causes the cooperators’ population to fluctuate around the critical population size. This significantly shortens the rescue time window during which cooperator mutants are able reach sufficient population size before the ancestral cooperator population drops below the critical size (**Fig. 5B**). Therefore the likelihood of evolutionary rescue is greatly reduced in the presence of cheaters. In fact, rescue only occured in our simulations in extreme cases, in which cooperators have a growth advantage of orders of magnitude over the cheaters when at low density (**Fig. 5A**).

**Figure 5:**
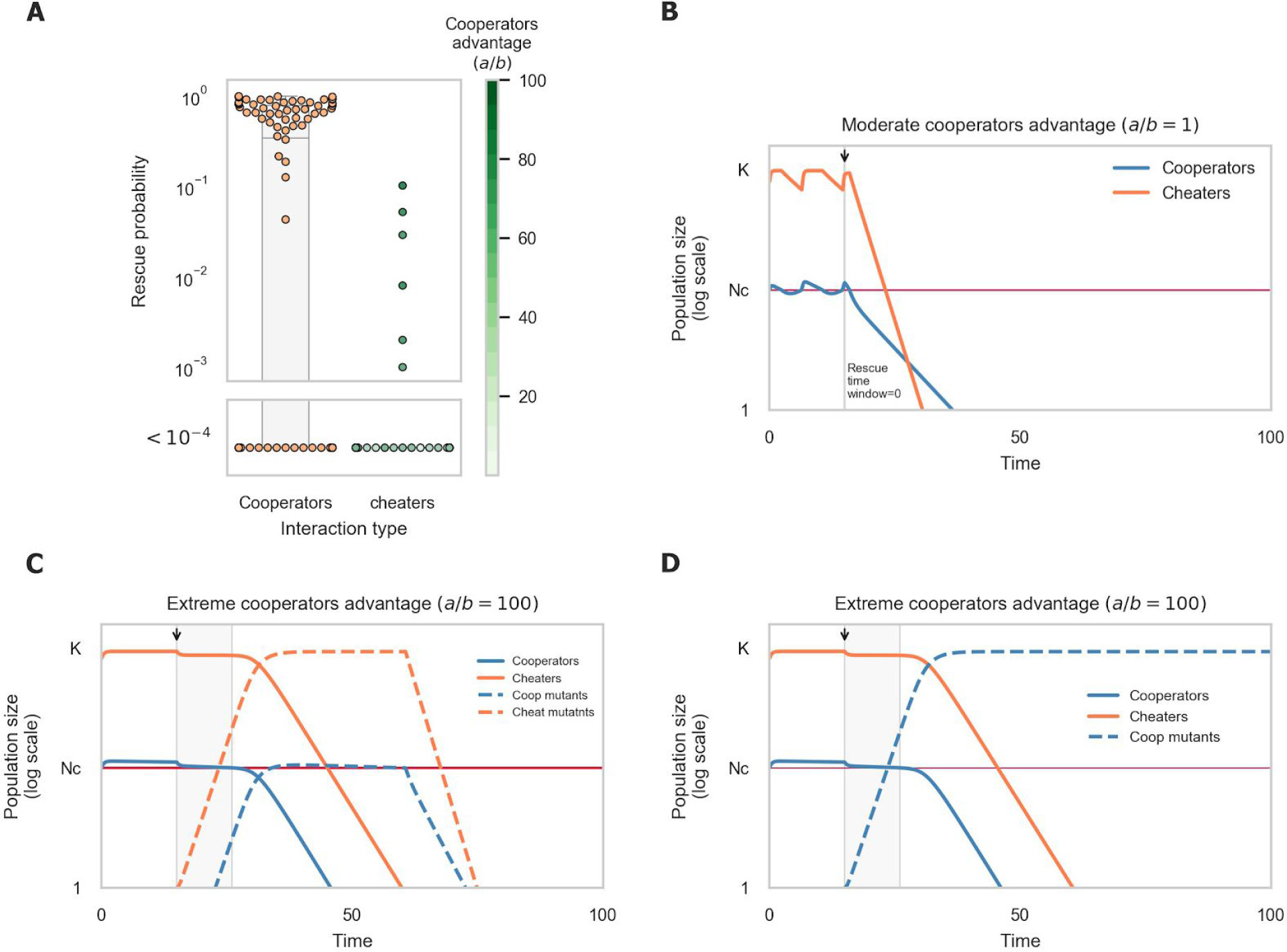
Evolutionary rescue of cooperative populations in the presence of cheaters is extremely unlikely. (A) Rescue probability of cooperative population in the presence of cheaters is orders of magnitude lower than with no cheaters. Dots represent the rescue probability calculated from simulations run with different sets of parameters (Critical population size (*N_c_*), growth rate of ancestor and mutant (*r_a_,r_m_*), cooperators advantage at high densities (*a*) and cheaters advantage when cooperator is at low density (*b*)). Rescue was observed only in extreme cases, in which cooperators have a growth advantage of orders of magnitude over the cheaters when at low density (B) Cooperators oscillate around critical population size prior to the onset of stress, eliminating the rescue time window. (C+D) Cheaters are purged from populations who manage to adapt to the new environments. When cheaters manage to adapt (C), cooperators mutants are pushed below critical populations size, causing both populations to collapse. Only populations in which cooperators adapt and cheaters do not survive (D).

In the extreme cases in which rescue occurs, cheaters are purged from the surviving population (**Fig. 5C-D**). When both cheaters and cooperators manage to adapt, cooperators are rapidly pushed below critical population size due to competition with the adapted cheaters, causing both populations to collapse. This resembles the competitive effect found in mutualisms: as a population with competitive advantage adapts to the environment, interference with other populations on which it is dependent can ultimately lead to collapse of the entire system.

Our results suggest strong group selection against populations invaded by cheaters in unstable environments. Adaptation is feasible only in populations in which the cooperators’ advantage in low densities is extremely high. In addition, the fact that evolutionary rescue requires that cheaters do not adapt to the new environment constitutes a selective pressure towards purging of cheaters from the population.

## Discussion

Our findings reveal that positive interactions can significantly decrease populations’ likelihood of evolutionary rescue (**Fig. 6A**). We found that this reduction is mainly due to the fact that survival in such populations requires at least a minimal number of cooperating individuals, reducing the time window during which adapted mutants can rise and spread. In mutualistic populations, we observed that the reduction of rescue probability is exacerbated by two additional effects: First, due to codependency between the two mutualistic partners, the rise and spread of adapted mutants in each of the populations is required in order to prevent either of the populations from collapsing. Second, due to competition for resources, adaptation of one of the species accelerates the decline of its partner towards extinction. Finally, we demonstrated that the presence of cheaters reduces the likelihood of evolutionary rescue even further, making it extremely unlikely, primarily since the cooperator population is close to its critical population size prior to the onset of the stress.

**Figure 6:**
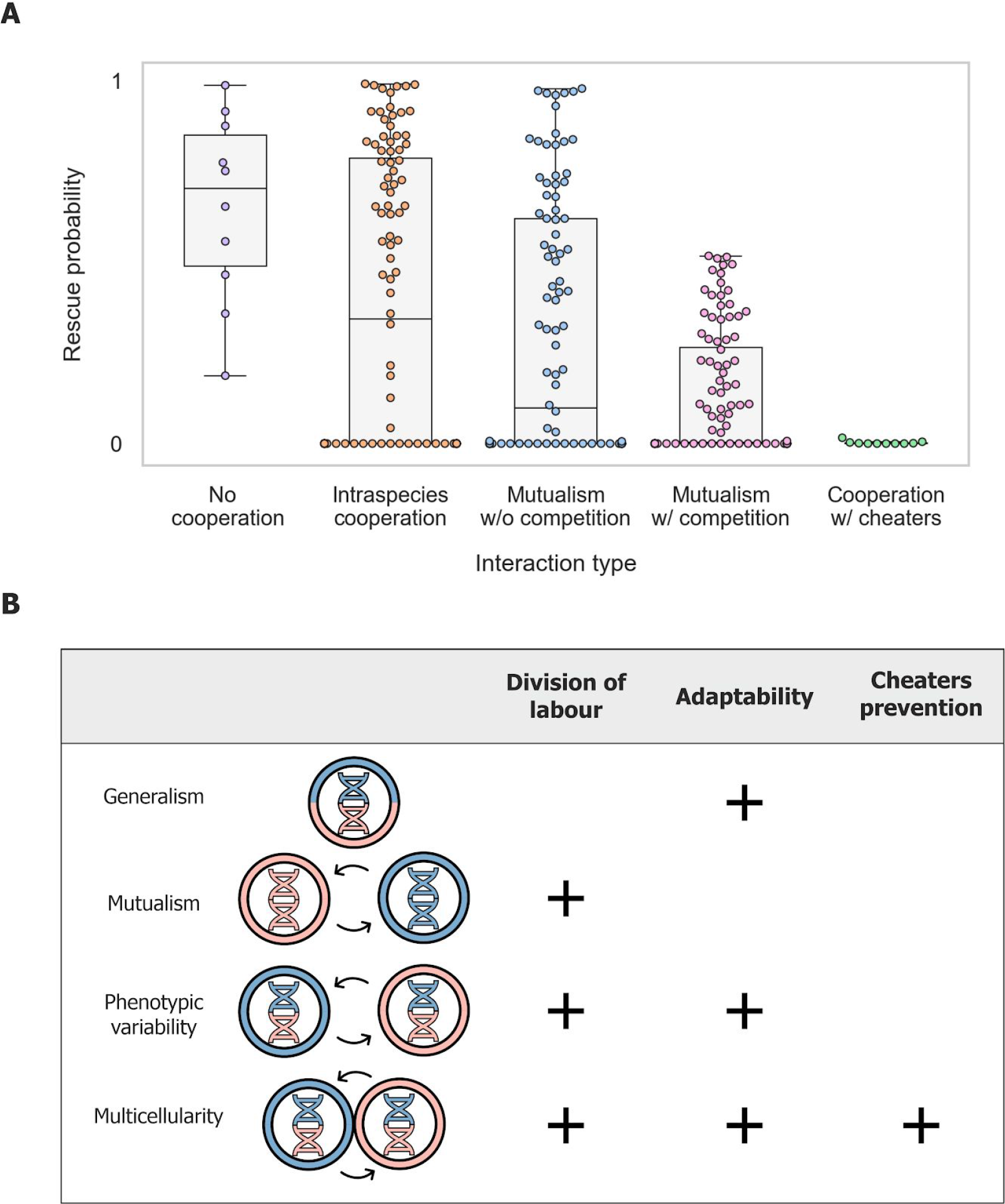
The reduced adaptability of positive interactions may have contributed to the formation of more complex strategies for division of labor. (A) Summary of observed rescue probability for different types of interactions. Dots represent the rescue probability that resulted from simulations run with different sets of parameters as in figure 2. (B) Different strategies of cooperation and the selective advantage they permit. The combined selection pressure imposed by the need to divide labour, adapt to novel environments and limit exploitation by cheaters may select for the formation of populations that differentiate into individuals with specialized phenotypes that form tight spatial structures, and potentially even multicellular organisms. Colors represent two different tasks required for growth, helix and circle colors represent genotype and phenotype, respectively.

Since populations engaged in positive interactions are more prone to collapse due to their reduced capacity for evolutionary rescue, how can they be so prevalent in nature? One possibility is that positive interactions occur primarily in steady environments, where frequent adaptation via evolutionary rescue is not essential. In addition, positive interactions may occur transiently - arising when the environment is stable due to selection for gene loss or division of labor, and collapsing when conditions change dramatically and adaptation is required. Lastly, positive interactions may be more stable when they occur as mutualism between species that do not compete strongly, since such competition greatly reduces mutualisms’ evolutionary rescue probability (**Fig. 6A**). For example, little or no competition occurs between plants and their mutualistic pollinator partners^42,43^.

In changing environments, where severe stresses occur periodically, there can be a conflict between maintaining the capacity for adaptation and gaining the benefits of division of labor (**Fig. 6B**). Mutualism offers the benefits of division of labor but has low adaptability. In contrast, generalism, where each individual performs all the tasks required for its growth, has high adaptability but does not provide the benefits of division of labor. A strategy that can offer the benefits of division of labour while maintaining the capacity for adaptation is phenotypic variability - a single genotype that differentiates into several specialized phenotypes^18,44,45^. In this situation, division of labour is enabled by positive interactions between different specialized phenotypes, while the capacity for adaptation is maintained since these phenotypes share a single genome. Thus both the interacting phenotypes can adapt via a single adaptive mutation, and the competitive effect that occurs during the adaptation of two mutualistic species is circumvented.

Our observation that the presence of cheaters significantly lowers populations’ ability to adapt underscores the importance of cheater avoidance mechanisms when facing environmental changes. Prevention of cheaters invasion is commonly achieved through population spatial structure, which enables cooperators to preferentially interact with each other^7,46–48^. Therefore, in changing environments, the combined selective pressure imposed by the need to divide labour, adapt to novel environments, and limit exploitation by cheaters may select for the formation of populations that differentiate into individuals with specialized phenotypes that form tight spatial structures (**Fig. 6B**). Strikingly, several organisms that are thought to be precursors to true multicellularity, such as choanoflagellates and volvox, fulfil these criteria^47–50^. More broadly, the elevated risk of extinction experienced by cooperative groups invaded by cheaters generates strong selection against cheaters at the group level^49–51^. Such group or lineage selection is considered a key factor leading to the evolution of multicellular organisms^52,53^. Thus, our insights into the influence of positive interactions on adaptation suggest new selective forces that might have been involved in the transition between unicellular and multicellular life.

Our results demonstrate the importance of considering ecological interactions when addressing evolutionary questions. Research interest in species adaptation to new environments has rapidly increased in recent years due to the realization that human activities are causing major changes to the environment of numerous ecosystems. Our findings highlight the potential of interactions within ecosystems to alter their fate in the face of environmental changes, such as those caused by anthropogenic influences.

## Methods

### Numerical Simulations

We have constructed simple models to describe the dynamics of positive interactions when evolutionary rescue is required, which is detailed in the supplementary information. Briefly, the model extends the logistic growth rate by applying strong Allee effect and addition of external growth rate (eq. 1–10 and supplement). Each simulation began with the growth of an ancestor population in an unstressed environment (*δ* = 0), followed by an onset of stress that increases the death rate such that it exceeds the ancestral exponential growth rate (*r_A_* < *δ*), leading the population to decline toward extinction. Simulations were ran in discrete time intervals (Δ*t* = 0.01), and the number of mutants *M* that arose during a time interval was sampled as a Poisson process, with the expected number of mutants determined by the ancestral population size and the mutation rate (eq. S4). Throughout the simulations, populations whose size decreased below 1 were considered to be extinct. In addition, to maintain simplicity, no further stochastic effects were considered in this model. Simulations end when extinction or rescue occurs, as defined for each model in the supplementary material. Evolutionary rescue probability was calculated by running 1000 simulations for each parameter set (Table S1), and calculating the fraction of simulations that resulted in rescue. The simulations were implemented using custom python scripts, using Scipy integrator for the calculation of population size change within each time step. The code is available upon request.

## Acknowledgments

We thank Nadav kashtan and Alfonso Pérez Escudero for constructive comments on the manuscript, and the members of the Friedman lab for helpful discussions.

## References

1. Boucher, D. H., James, S. & Keeler, K. H. The Ecology of Mutualism. Annu. Rev. Ecol. Syst. 13, 315–347 (1982).

2. Bronstein, J. L. Our Current Understanding of Mutualism. Q. Rev. Biol. 69, 31–51 (1994).

3. Bruno, J. F., Stachowicz, J. J. & Bertness, M. D. Inclusion of facilitation into ecological theory. Trends Ecol. Evol. 18, 119–125 (2003).

4. Callaway, R. M. Positive interactions among plants. Bot. Rev. 61, 306–349 (1995).

5. Bruno, J. F. & Bertness, M. D. Habitat modification and facilitation in benthic marine communities. (2001).

6. Clutton-Brock, T. Breeding Together: Kin Selection and Mutualism in Cooperative Vertebrates. Science 296, 69–72 (2002).

7. West, S. A., Griffin, A. S., Gardner, A. & Diggle, S. P. Social evolution theory for microorganisms. Nat. Rev. Microbiol. 4, 597–607 (2006).

8. Faust, K. et al. Microbial Co-occurrence Relationships in the Human Microbiome. PLOS Comput. Biol. 8, e1002606 (2012).

9. Mainali, K., Bewick, S., Vecchio-Pagan, B., Karig, D. & Fagan, W. F. Detecting interaction networks in the human microbiome with conditional Granger causality. PLOS Comput. Biol. 15, e1007037 (2019).

10. Dugatkin, L. A., Perlin, M. & Atlas, R. The Evolution of Group-beneficial Traits in the Absence of Between-group Selection. J. Theor. Biol. 220, 67–74 (2003).

11. Brook, I. Beta-lactamase-producing bacteria in mixed infections. Clin. Microbiol. Infect. Off. Publ. Eur. Soc. Clin. Microbiol. Infect. Dis. 10, 777–784 (2004).

12. Mariscal, R. N., Fautin, D. G. & Allen, G. R. Field Guide to Anemonefishes and Their Host Sea Anemones. Copeia 1993, 899 (1993).

13. Morris, J. J., Lenski, R. E. & Zinser, E. R. The Black Queen Hypothesis: Evolution of Dependencies through Adaptive Gene Loss. mBio 3, e00036–12 (2012).

14. Morris, J. J., Papoulis, S. E. & Lenski, R. E. Coexistence of Evolving Bacteria Stabilized by a Shared Black Queen Function. Evolution 68, 2960–2971 (2014).

15. Shapiro, J. A. Thinking about bacterial populations as multicellular organisms. Annu. Rev. Microbiol. 52, 81–104 (1998).

16. Gao, Y., Traulsen, A. & Pichugin, Y. Interacting cells driving the evolution of multicellular life cycles. PLOS Comput. Biol. 15, e1006987 (2019).

17. Michod, R. E. & Roze, D. Cooperation and conflict in the evolution of multicellularity. Heredity 86, 1–7 (2001).

18. Wahl, L. M. The Division of Labor: Genotypic versus Phenotypic Specialization. Am. Nat. 160, 135–145 (2002).

19. Wolk, C. P., Ernst, A. & Elhai, J. Heterocyst Metabolism and Development. in The Molecular Biology of Cyanobacteria (ed. Bryant, D. A.) 769–823 (Springer Netherlands, 1994). doi:10.1007/978-94-011-0227-8_27.

20. Dunn, R. R., Harris, N. C., Colwell, R. K., Koh, L. P. & Sodhi, N. S. The sixth mass coextinction: are most endangered species parasites and mutualists? Proc. R. Soc. B Biol. Sci. 276, 3037–3045 (2009).

21. Rezende, E. L., Lavabre, J. E., Guimarães, P. R., Jordano, P. & Bascompte, J. Non-random coextinctions in phylogenetically structured mutualistic networks. Nature 448, 925–928 (2007).

22. Memmott, J., Craze, P. G., Waser, N. M. & Price, M. V. Global warming and the disruption of plant-pollinator interactions. Ecol. Lett. 10, 710–717 (2007).

23. Hardin, G. The Tragedy of the Commons. Science 162, 1243–1248 (1968).

24. Rankin, D. J., Bargum, K. & Kokko, H. The tragedy of the commons in evolutionary biology. Trends Ecol. Evol. 22, 643–651 (2007).

25. Frost, I. et al. Cooperation, competition and antibiotic resistance in bacterial colonies. ISME J. 12, 1582–1593 (2018).

26. Griffin, A. S., West, S. A. & Buckling, A. Cooperation and competition in pathogenic bacteria. Nature 430, 1024–1027 (2004).

27. Hoek, T. A. et al. Resource Availability Modulates the Cooperative and Competitive Nature of a Microbial Cross-Feeding Mutualism. PLoS Biol. 14, (2016).

28. Machado, D. et al. Polarization of microbial communities between competitive and cooperative metabolism. bioRxiv2020.01.28.922583 (2020) doi:10.1101/2020.01.28.922583.

29. Abrams. Modelling the adaptive dynamics of traits involved in inter- and intraspecific interactions: An assessment of three methods. Ecol. Lett. 4, 166–175 (2001).

30. Foster, K. R. & Wenseleers, T. A general model for the evolution of mutualisms. J. Evol. Biol. 19, 1283–1293 (2006).

31. Frederickson, M. E. Rethinking Mutualism Stability: Cheaters and the Evolution of Sanctions. Q. Rev. Biol. 88, 269–295 (2013).

32. Litsios, G. et al. Mutualism with sea anemones triggered the adaptive radiation of clownfishes. BMC Evol. Biol. 12, 212 (2012).

33. Miller-Struttmann, N. E. et al. Functional mismatch in a bumble bee pollination mutualism under climate change. Science 349, 1541–1544 (2015).

34. Hillesland, K. L. & Stahl, D. A. Rapid evolution of stability and productivity at the origin of a microbial mutualism. Proc. Natl. Acad. Sci. 107, 2124–2129 (2010).

35. Yurtsev, E. A., Conwill, A. & Gore, J. Oscillatory dynamics in a bacterial cross-protection mutualism. Proc. Natl. Acad. Sci. 113, 6236–6241 (2016).

36. Adamowicz, E. M. & Harcombe, W. R. Weakest link dynamics predict apparent antibiotic interactions in a model cross-feeding community. bioRxiv 2020.03.10.986695 (2020) doi:10.1101/2020.03.10.986695.

37. Gomulkiewicz, R. & Holt, R. D. When does Evolution by Natural Selection Prevent Extinction? Evolution 49, 201–207 (1995).

38. Bell, G. & Gonzalez, A. Evolutionary rescue can prevent extinction following environmental change. Ecol. Lett. 12, 942–948 (2009).

39. Allee, W. C. & Bowen, E. S. Studies in animal aggregations: Mass protection against colloidal silver among goldfishes. J. Exp. Zool. 61, 185–207 (1932).

40. Kanarek, A. R. & Webb, C. T. ORIGINAL ARTICLE: Allee effects, adaptive evolution, and invasion success. Evol. Appl. 3, 122–135 (2010).

41. Sanchez, A. & Gore, J. Feedback between Population and Evolutionary Dynamics Determines the Fate of Social Microbial Populations. PLoS Biol. (2013) doi:10.1371/journal.pbio.1001547.

42. Waser, N. M. & Ollerton, J. Plant-Pollinator Interactions: From Specialization to Generalization. (University of Chicago Press, 2006).

43. Mutualism. (Oxford University Press, 2015).

44. Gestel, J. V., Vlamakis, H. & Kolter, R. Division of Labor in Biofilms: the Ecology of Cell Differentiation. in Microbial Biofilms 67–97 (John Wiley & Sons, Ltd, 2015). doi:10.1128/9781555817466.ch4.

45. Wloch-Salamon, D. M., Fisher, R. M. & Regenberg, B. Division of labour in the yeast: Saccharomyces cerevisiae. Yeast 34, 399–406 (2017).

46. Nowak, M. A. & May, R. M. Evolutionary games and spatial chaos. Nature 359, 826–829 (1992).

47. Gilbert, O. M., Foster, K. R., Mehdiabadi, N. J., Strassmann, J. E. & Queller, D. C. High relatedness maintains multicellular cooperation in a social amoeba by controlling cheater mutants. Proc. Natl. Acad. Sci. 104, 8913–8917 (2007).

48. Marchal, M. et al. A passive mutualistic interaction promotes the evolution of spatial structure within microbial populations. BMC Evol. Biol. 17, 106 (2017).

49. Alexander, R. D. & Bargia, G. Group Selection, Altruism, and the Levels of Organization of Life. Annu. Rev. Ecol. Syst. 9, 449–474 (1978).

50. Keller, L. Levels of Selection in Evolution. (Princeton University Press, 1999).

51. Dobata, S. & Tsuji, K. A cheater lineage in a social insect. Commun. Integr. Biol. 2, 67–70 (2009).

52. Griesemer, J. The Units of Evolutionary Transition. Selection 1, 67–80 (2005).

53. Rainey, P. B. Unity from conflict. Nature 446, 616–616 (2007).

